## Supplementary material for "Dual-specific autophosphorylation of kinase IKK2 enables phosphorylation of substrate IκBα through a phosphoenzyme intermediate": Materials and methods

**Reagents and Cell culture materials:**

Monoclonal mouse anti-IκBα (L35A5) (catalogue number: 4814, RRID: AB_390781), mouse anti-Phospho-IκBα (Ser^32/36^) (5A5) (catalogue number:9246, RRID: AB_2267145) and rabbit anti-Phospho-IKKα/β (Ser^176/180^) (16A6) (catalogue number: 2697, RRID: AB_2079382) were purchased from Cell Signaling Technology (CST). Monoclonal mouse anti-IKKβ (10AG2) (catalogue number: NB100-56509) was purchased from Novus Biologicals. Polyclonal rabbit anti-IKKβ (catalogue number: BB-AB0094) and Polyclonal rabbit anti-6XHis (catalogue number: BB-AB0010) were procured from Bio Bharti Life Science. Monoclonal mouse anti-Phospho-Tyrosine (catalogue number: 610000, RRID: AB_397423) was purchased from BD Biosciences or from EMD Millipore (catalogue no. 05-321X). Polyclonal rabbit anti-GST (catalogue number: 924801, RRID: AB_2565461), monoclonal mouse anti-HA 11 epitope (catalogue number: 901502, RRID: AB_2565007) and mouse anti-Tubulinβ3 (catalogue number: 657402, RRID: AB_2562570) were purchased from BioLegend. Polyclonal rabbit anti-IKKα/β (H-470) (catalogue number: sc7607, RRID: AB_675667) was purchased from Santa Cruz Biotechnology (has been discontinued) and mouse anti-Phospho-Serine Q5 (catalogue number: 37430) was from Qiagen. Horseradish peroxidase-conjugated secondary antibodies against rabbit (catalogue number: 31460, RRID: AB_228341) and mouse IgG (catalogue number: 31430, RRID: AB_228307) were Pierce antibodies purchased from Thermo Scientific. Adenosine-5'-Triphosphate (ATP), Disodium trihydrate (catalogue number: A-081-25) was obtained from Gold Biotechnology and Adenosine Diphosphate (ADP) (Reference number: V916A) used was provided with ADP-Glo^TM^ Kinase assay kit (catalogue number: V6930) from Promega. Recombinant murine TNF-α used was obtained from Roche.

Sf9 cells (Gibco™ Sf-900™ III SFM) for expression of proteins using baculovirus system was procured from Thermo Scientific (catalogue number: 12659017) and were maintained in media Gibco™Sf-900™ III SFM, Thermo Scientific (catalogue number: 12658019). Cells were grown and maintained as per the manufacturer’s protocol.

Immortalized MEF cells were grown in DMEM supplemented with 10% heat-inactivated bovine calf serum, penicillin (100units/ml), streptomycin (100μg/ml) and L-glutamine (1%) at 5% CO_2_ and 37°C. Authenticity of MEF cells were carefully assessed by observing their characteristic spindle shape morphology and their conformity to 3T3 fibroblasts, and by observing contact inhibition properties. The cells were also routinely assessed for the presence/absence of IKK1 and IKK2, and IκBα degradation dynamics in response to mouse TNFα treatment using immunoblotting (Polley et al., 2016). Mycoplasma contamination/infection leads to the degradation of IκBα protein and NF-κB activation even in absence of any known external signaling cues (i.e., control untreated cells) and thus easily recognized and could be further tested. In summary, cells were carefully maintained as adherent monolayer cultures using the 3T3 protocol, and cells not conforming to ideal phenotypic or functional NF-κB signaling characteristics were discarded.

**Protein expression and purification:**

**IKK2 (WT and other constructs):**

Wild-type and mutant IKK2 proteins were purified following a published protocol (Polley et al, 2013). Sf9 cells (Gibco™ Sf9 cells in Sf-900™ III SFM) at a density of 1.5-2 million/ml were infected with a high titre viral stock (generally stage P3 or beyond) at a ratio (usually around 1:100) optimized for a high expression level through a pilot experiment, and cultured for 48-60 hours upon infection. Cells were harvested and lysed by sonication in a buffer containing 25 mM Tris-HCl pH 8.0, 200 mM NaCl, 10% Glycerol, 5 mM β-Mercaptoethanol (β-ME), 10 mM Imidazole, 20 mM β-glycerophosphate, 10 mM NaF, 1 mM sodium orthovanadate, and 1X Protease Inhibitor Cocktail (PIC, Sigma). The lysate was clarified by centrifugation at 20,000 x g for 45 minutes at 4˚C and incubated with Ni-NTA agarose resin (Qiagen, catalogue number: 30230) pre-equilibrated with lysis buffer for 2 hours at 4˚C. The resin was subsequently washed with lysis buffer containing 30 mM Imidazole. Protein was eluted with lysis buffer containing 250 mM Imidazole. Protein fractions were analysed on SDS-PAGE, pooled together, concentrated, and either flash frozen or subjected to size-exclusion chromatography on Superdex 200 Increase 10/300 GL (Cytiva) column equilibrated with 20 mM Tris-HCl pH 8.0/ 25 mM HEPES pH 7.9, 200 mM NaCl, 10% Glycerol and 2 mM DTT. To produce untagged IKK2, protein fractions eluted from the Ni-NTA resin of desired purity was treated with TEV-protease at a ratio of 50:1 (weight by weight, IKK2: TEV) at 4˚C for 10 hours or longer in presence of 1 mM of EDTA, the resulting protein mix was then loaded on to the SEC as described above. To prepare autophosphorylated IKK2, the fractions with desired purity level were treated with 0.5-1 mM of ATP, 10 mM of MgCl_2_ and phosphatase inhibitors and the reaction mixture was incubated at 27˚C for 1 hour. Resulting reaction mixture was concentrated in a YM30-concentrator and loaded onto a Superdex 200 Increase 10/300 GL column. Elution fractions of the size-exclusion chromatography were analysed on an SDS-PAGE, and peak fractions of desired quality were pooled together and concentrated using a YM30 Centriprep (Millipore) or Centricon devices (Millipore) depending upon the volume of the sample, and flash frozen in liquid nitrogen immediately.

**IκBα (WT, mutant and other constructs):**

His-tagged full-length and GST-tagged deletion (1-54) constructs of IκBα (both the wild-type and Ser32Ala, Ser36Ala, Ser32Glu, Ser36Glu, Ser32Ala,Ser36Ala single and double-mutant versions) were expressed in *E. coli* Rosetta2 (DE3). The cells at an attenuance of 0.6 were induced with 0.3 mM IPTG and cultured overnight at 16˚C. The cell pellet was resuspended in 40 ml of Lysis Buffer (25 mM Tris pH 7.5, 200 mM NaCl, 10 mM imidazole, 10% Glycerol, 5 mM β-mercaptoethanol, and 1 mM PMSF). The cells were lysed on ice with a sonicator. The lysate was clarified by centrifugation at 15,000 x g for one hour at 4°C and mixed with 2-3 ml slurry of Ni-NTA Agarose resin that was equilibrated with lysis buffer on a rotary mixer for 2 hours in a 4°C room. The resin was washed with the lysis buffer containing 500 mM NaCl (high salt wash) for at least 50 column volume to remove non-specifically bound proteins. Subsequently, the resin was thoroughly washed with the lysis buffer containing 20 mM imidazole. His-tagged IκBα protein was eluted under gravity flow, using elution buffer (the lysis buffer containing 250 mM imidazole), and elution fractions of ~1 ml volume were collected. For GST tagged proteins, Glutathione Sepharose resin (Cytiva, catalogue number: 17-5279-01) was used, and the elution was performed with the help of reduced glutathione). The elution fractions containing quality proteins were subjected to size-exclusion chromatography (SEC) on a HiLoad 16/600 pg Superdex 200 column (Cytiva) in 25 mM Tris pH 7.5, 100 mM NaCl, 5% glycerol and 5 mM DTT. The peak fractions were concentrated, aliquoted, and stored at -80˚C.

**NEMO FL WT:**

His-tagged FL NEMO WT was expressed in *E. coli* Rosetta2 DE3 cells. The cells were grown at 37˚C in LB medium containing 50 μM ZnCl_2_ to an attenuance of 0.3-0.4 and(Millipore) cooled down to 20 ˚C. Upon addition of 0.3 mM IPTG, the culture was further incubated for 14-16 hours at 20˚C. The culture post-induction was cooled down to 4˚C before harvesting cells by centrifugation at 5000 rpm at 4˚C. The cell pellet was resuspended in 40 ml (per litre of culture) of Lysis Buffer (25 mM Tris pH 7.5, 200 mM NaCl, 10 mM imidazole, 10% Glycerol, 5 mM β-mercaptoethanol, 20 μM ZnCl_2_, 0.5 M urea (as an osmolyte). The rest of the steps performed were similar to purification with Ni-NTA described above. The main elution fractions were subjected to size-exclusion chromatography on a HiLoad 16/600 pg Superdex 200 column (Cytiva) in 25 mM HEPES, 100 mM NaCl, 5% glycerol and 5 mM β-mercaptoethanol buffer. The cleaner peak fractions were stored pooled, concentrated, and stored at -80˚C.

**Kinase Assay with purified proteins:**

The presented kinase assay figures are ideal representatives of a minimum of 3 similar experiments, unless otherwise noted. For all *in vitro* kinase assays, a master mix was prepared just prior to reaction start to minimize pipetting and other errors. Typically, 50-100 ng (unless otherwise noted) of purified kinase was used in each kinase reaction of 20μl volume. 0.5-1μg of IκBα substrate (full-length or GST-tagged 1-54) was incubated with the kinase for 30 mins or indicated time periods at 27˚C in presence of 20μM ATP spiked with 0.1μCi of γ-P^32^-ATP in case of radioactive assay or 50-100 μM ATP in case of Western blot assays in a reaction buffer containing 20mM HEPES pH 7.5, 100mM NaCl, 15mM MgCl_2_, 2mM DTT, 0.2mM Na_3_VO_4_, 10mM NaF, 20mM β-Glycerophosphate. Reaction was stopped by addition of 1X-Laemmli buffer and heating the samples to 95˚C for 5 mins. Samples (half of the total reaction mixture) were resolved on 8 or 10% SDS-PAGE gels. For radioactive assays, the gels were quickly stained and de-stained by Coomassie staining protocol to confirm loading equivalencies. Typically, the substrate band(s) were monitored for this purpose as kinase band(s) were not visible in quick Coomassie staining protocol. For radioactive kinase assays with IKK2 K44M mutant, a higher amount of kinase was used (0.5-1μg), and the kinase band was visible following the same staining/de-staining protocol (Fig. 3C and S3E, Fig. S1A). The gels were subsequently dried and analysed using Typhoon phosphor imager or by exposing it to Kodak films. For Western blot analyses, proteins from the SDS-PAGE were electro-transferred onto nitrocellulose/PVDF membrane and probed with desired antibodies. Phosphorylated tyrosine and serine residues were detected using monoclonal antibodies against phospho-tyrosine and phospho-serine. Specific phosphorylation on S32/S36 of IκBα was detected using a monoclonal antibody against IκBα phosphorylated at S32/36, and phosphorylation at the activation loop serines of IKK2 was detected using a monoclonal antibody against IKK2 phosphorylated at S177/S181.

For the assays with inhibitors, IKK2 or IKK2:IκBα were mixed with the respective inhibitors in the kinase assay buffer at concentrations indicated in the respective figures for 30 minutes at room temperature - prior to the addition of ATP or γ-P^32^-ATP. Upon addition of ATP, reactions were allowed to proceed for indicated time periods prior to analysis by autoradiography or immunoblotting as described above.

**IP-Kinase assay:**

Immortalized *ikk2*^-/-^ mouse embryonic fibroblast 3T3 cells were reconstituted by retroviral transduction with HA-tagged WT or mutant pBABE-IKK2 constructs. Stable expression of IKK2 was confirmed by immunoblotting. Freshly plated cells were stimulated with recombinant murine TNF-α (Roche) when cells reached confluency of ~ 70-80%. Whole cell extract was prepared and pre-cleared by incubation with Protein A agarose bead for an hour. IKK complex was immunoprecipitated from whole cell extracts using a NEMO-specific antibody (BD) and Protein A agarose (Upstate) for 2 hr at 4˚C. Immunoprecipitated complex was washed extensively, including a final washing step with the kinase assay buffer devoid of ATP and incubated with GST-IκBα (1-54) and 20 μM ATP spiked with 5 μCi of γ-^32^P-ATP for 30 mins at 30˚C in kinase assays. Reactions were stopped by addition of 2X-Laemmli SDS buffer and heating at 95˚C for 5 min prior to resolving on a 12% SDS PAGE. Extent of substrate phosphorylation was measured by cutting out the appropriate area of the gel containing the substrate and exposing it to phosphor-imager plates (Typhoon, Cytiva) after drying. For control of the amount of IKK2 in each reaction, IKK2 was transferred from gel onto nitrocellulose membrane and probed with anti-IKK1/2 antibody. These assays were performed twice and representative data images are shown.

**Sample preparation for LC-MS/MS analyses for Tyrosine phosphorylation detection:**

A kinase assay reaction was set up with about 20 µg of FL IKK2 WT in the presence of 100 µM ATP for about an hour at 27°C. SDS to a final concentration of 0.1% was added and reaction was incubated at 30°C for 30 minutes. The sample buffer was exchanged to 50 mM Ammonium bicarbonate (Sigma), 8 M urea (Merck) and 2.5 mM TCEP using a 10 kDa concentrator (EMD Millipore), and sample was further incubated at 37°C for 45 minutes. Next, 20 mM iodoacetamide (IAA) was added (final concentration) to the sample and incubated in the dark at room temperature for an hour. After this step, 5 mM Dithiothreitol (DTT) (Promega, MS grade) was added and incubated at room temperature for an hour. The sample was then passed through a protein-desalting column PD SpinTrap (GE Healthcare) to get rid of urea. GluC protease in the final ratio of protease: protein i.e. 1:50 (w/w) was then added and incubated for an hour at 37°C. Next, Trypsin Gold (Promega, MS grade) was added to the sample at a final ratio of protease: protein i.e. 1:100 (w/w) and incubated at 37°C for 2 hours. Finally, Trypsin Gold was added in the final ratio of 1:50 (w/w) to the sample for overnight incubation at 37°C. On the following day, 0.1% of formic acid (final concentration) was added to inactivate trypsin and incubated for 5 minutes at room temperature. The samples were quickly frozen at -80°C and then dried out using a vacuum evaporator (Savant RVT5105, Thermo Scientific). The trypsinized sample was then resuspended in 0.5% TFA and 5% ACN solution. The sample was passed through the C18 spin column (Pierce, Thermo Scientific) by following the manual. Finally, the elution was performed in 70% ACN and 0.1% formic acid and then elute was dried out using the vacuum evaporator.

**Preparation of FL IKK2 K44M for mass spectrometry to check purity**

About 25 µg of purified protein was taken and denatured at 90°C for 10 minutes. 3 mM Tris (2-carboxyethyl) phosphine (TCEP) (GoldBiochem) was then added and reaction was incubated for 45 minutes at 37°C. Next, 20 mM iodoacetamide (IAA) was added (final concentration) to the sample tube and incubated in the dark at room temperature for an hour. After this step, 5 mM Dithiothreitol (DTT) (Promega, MS grade) was added and incubated at room temperature for an hour to neutralise the effect of excess IAA. Next, Trypsin Gold (Promega, MS grade) was added to the sample at a final ratio of protease: protein i.e. 1:100 (w/w) and incubated at 37°C for 2 hours. Finally, Trypsin Gold was added in the final ratio of 1:50 (w/w) to the sample for overnight incubation at 37°C. On the following day, 0.1% of formic acid (final concentration) was added to inactivate trypsin and incubated for 5 minutes at room temperature. The samples were quickly frozen at -80°C and then dried out using a vacuum evaporator (Savant RVT5105, Thermo Scientific). The trypsinized sample was then resuspended in 0.5% TFA and 5% ACN solution. The sample was passed through the C18 spin column (Pierce, Thermo Scientific) following the manufacturer’s protocol. Finally, the elution was performed in 70% ACN and 0.1% formic acid and elute was dried out using the vacuum evaporator.

**LC-MS/MS analysis:**

For analyses on an Orbitrap platform, peptides were resuspended in 5% (v/v) formic acid and sonicated for 5 min. Samples were analyzed on Orbitrap Exploris^TM^ 240 mass spectrometer (Thermo Scientific) coupled to a nanoflow LC system (Easy nLC II, Thermo Scientific). Peptides were loaded onto a PepMap^TM^ RSLC C18 nanocapillary reverse phase HPLC column (75 µm × 15 cm; 3μm; 100 Å) and separated using a 60 min linear gradient of the organic mobile phase [5% Acetonitrile (ACN) containing 0.2% formic acid and 90% ACN containing 0.2% formic acid]. For identification of peptides the raw data was analysed on MaxQuant proteomics computational platform (Ver. 1.6.8) (PMID:19029910) and searched against UniProt amino acid sequences (Uniprot Reference Proteome ID: UP000829999; Uniprot ID for IKK2: O14290). MaxQuant used a decoy version of the specified database to adjust the false discovery rates for proteins and peptides below 1%. The search parameters included constant modification of cysteine by carbamidomethylation, phosphorylation (STY) as a variable modification and enzyme specificity as trypsin. iBAQ option was selected to compute abundance of the proteins.

**ESI-MS analyses of ADP:**

A 50 μM solution of ADP was prepared for analysis by Electrospray Ionisation Mass Spectrometry (ESI-MS) in a Xevo G2-XS QToF mass spectrometer (Waters Corporation) using capillary at 3kV, with voltage parameters of sampling cone at 40V and source offset at 80V. The source temperature was set at 100˚C and desolvation temperature at 40˚C. The gas flow rates were as follows: Cone gas flow rate at 50 litre/hr and desolvation gas flow rate at 400 litre/hr. The sample was passed at 5 μl/min flow rate. The MS data was acquired for a mass range of 50-2000 m/z and the acquisition time was 1 minute.

**Radioactive Phospho-transfer assay:**

Phospho-transfer assays were performed in a wide range of concentration of IKK2, IκBα, and ATP. Typically, 10 μM concentration of purified IKK2 protein was autophosphorylated in presence of 100-200 nM of γ^32^P-ATP (PerkinElmer) in kinase assay buffer for 1 hour at 30˚C. 70 μl of the reaction mixture was passed through a Micro Bio-Spin Bio-Gel P-30 spin column (Bio-Rad; 40 kDa MWCO) equilibrated with 20 mM HEPES pH 7.9, 100 mM NaCl, 0.2 mM Na_3_VO_4_, 10 mM NaF, 20 mM β-Glycerophosphate, 2 mM DTT, 2 mM EDTA and 1X protease inhibitor cocktail. Eluted protein was subjected to another round of buffer exchange on a Bio-Gel P-30 column equilibrated with the same buffer mentioned above devoid of EDTA and containing 15 mM MgCl_2_. Phosphorylated IKK2 thus obtained was used in phosphotransfer reaction. 2-3 μM of Phosphorylated IKK2 was incubated with 10-20 μM of WT or mutant IκBα protein at 30˚C in kinase assay buffer in a total volume of 20 μl for different time periods as indicated in Figure 5A and 6A. Presented figure is a representative of a minimum of 3 similar assays unless otherwise noted. This assay was performed twice with the mutant IκBα protein (Fig. 6A). The reactions were stopped by addition of 10 μl of 4X Laemmli SDS-buffer containing 100 mM EDTA to samples and heating at 95˚C for 5 min. It was observed that addition of SDS and EDTA was sufficient to stop the reaction without heating. Phopsho-transfer signal was not dependent upon heating of the reaction mixture. Different concentration of ADP or ATP was added to the assay to assess the role of these nucleotides in phosphotransfer efficiency. Routinely, 10-15μl of reaction products were resolved on 10% SDS PAGE and radiolabelled samples were quantified by exposing the dried gel to phosphor-imager plates scanned on a Typhoon scanner. For the phosphotransfer reaction described in Figure 5A, 10 μM purified IKK2 was autophosphorylated with 100 nM of γ^32^P-ATP in presence of 40 μM of cold ATP. Each concentration mentioned above is final concentration.

**Phospho-transfer assay with cold ATP:**

An ~ 10 μM of purified FL IKK2WT was mixed with 100 μM of cold ATP in 600 μl volume of kinase assay buffer (containing 1X protease inhibitor cocktail) for one hour at 30˚C. From the resulting reaction mixture, 500 μl was loaded onto a Superdex 200 increase 10/300 GL (24ml bed volume, Cytiva) SEC column to separate autophosphorylated FL IKK2WT from excess ATP. SEC was performed in buffer containing 20 mM HEPES, 100 mM NaCl, 20 mM β-glycerophosphate, 10 mM NaF, 0.2 mM Na_3_VO_4_ and 2 mM DTT. The elution fractions containing phosphorylated IKK2 were incubated with FL IκBαWT in presence of different concentrations of ATP or ADP at different time points (as mentioned in Figure 5E) at 30˚C. The reactions were stopped by addition of 4X Laemmli SDS dye and heating at 95˚C for 5 min and prior to resolving on a 10% SDS-PAGE. Immunoblotting was performed using monoclonal antibodies against phospho-tyrosine and phospho Ser32/36 IκBα to detect phosphorylations. This assay was performed twice and the representative figure is shown (Figure 5E).

**Molecular Dynamics Simulation**

The kinase domain structure of IKK2 (PDB ID: 4E3C), spanning residues 16 to 310, was obtained from the PDB database (Berman et al., 2000). We created unphosphorylated forms (at Ser177 & Ser181) and two phosphorylated forms (phosphorylated at Ser177 & Ser181 and at Tyr169 & Ser177 & Ser181) of IKK2 using UCSF CHIMERA software (Pettersen et al., 2004). These structures underwent molecular dynamics (MD) simulations using the GROMACSv2023.1 simulation package (Abraham et al., 2015), enabling a comparative analysis of their structural and energetic stability. The Charmm36 force field was applied to generate coordinates and topology files for the IKK2 structures (Huang and MacKerell, 2013). For simulation, a cubic box was defined and filled with TIP3 water molecules (Jorgensen et al., 1983). Sodium and chloride ions were added to neutralize the three systems. The system underwent two-stage minimization using steepest-descent (Nocedal and Wright, 1999) and conjugate-gradient algorithms (Straeter, 1971). Equilibration was carried out under NVT (constant particle number, volume, and temperature) and NPT (constant particle number, pressure, and temperature) conditions for 200 ps at 300 K and 1 atm pressure. MD simulations were conducted for 200 ns for each protein. To assess global structural changes, Root Mean Square Deviation (RMSD) analysis was employed, while Root Mean Square Fluctuation (RMSF) analysis revealed the overall and local motion/flexibility of the proteins. The *gmx_energy* module was utilized to compute energy during simulation, and *gmx_sasa* was employed to determine the solvent accessible surface area (SASA) of the protein. To understand the correlations of the atomic motions in different regions, the dynamic cross-correlation analysis was carried out. The initial step of dynamic cross-correlation analysis was to construct the covariance matrix which examines the linear relationship of atomic fluctuations for individual atomic pairs. The covariance matrix was calculated using the *gmx_covar* tool. Dynamic cross-correlation matrices (DCCM) were calculated from the C_α_-trace covariance matrix principal components. Results are displayed as a color coded with a value of −1 indicated completely anti-correlated motions and a value of +1 indicating completely correlated motions.

**Molecular Docking Analysis**

The initial (0^th^ ns) and terminal (200^th^ ns) structures from the simulations of unphosphorylated (UnP-IKK2), double phosphorylated (p-IKK2), and triple phosphorylated (P-IKK2) forms were subjected to molecular docking analysis using ATP and ADP as ligands, allowing for comparison of binding affinities. LeDock (Liu and Xu, 2019) and GOLD software were employed for ligand-protein docking. IKK2: ATP docking poses were compared with available kinase:ATP complexes, e.g., PKA: ATP complex (PDB ID: 1ATP) by calculating the ligand root mean square deviation (RMSD). The pose that resembled most with the pose found in PDB ID 1ATP were chosen for further analysis. Similarly, IKK2: ADP docking poses were also compared, where IRE1: ADP (PDB ID: 2RIO) complex was chosen for comparison and the pose closest to that in PDB ID: 2RIO were chosen for further analysis. Finally, the best selected poses were rescored using AutoDock Vina to represent the predicted binding affinities.

For assessing the ability of P-IKK2 to accommodate ATP or ADP, the MD-simulation-derived P-IKK2 structure was superimposed on the p-IKK2 structure bound to ATP/ADP, and compared as described.

**Estimation of Binding Free Energy**

Following the identification of the best docking pose, another round of molecular dynamics (MD) simulation was conducted using GROMACS for 10ns. 50 intermediate complex structures were extracted from each simulation run. The binding free energy (ΔG) of these intermediate structures was calculated using the MM-PBSA (Molecular Mechanics Poisson-Boltzmann Surface Area) method.
