## Supplementary figures and images for "Dual-specific autophosphorylation of kinase IKK2 enables phosphorylation of substrate IκBα through a phosphoenzyme intermediate"

### Supplementary Figure 1

#### Figure 1 - Figure Supplement 1

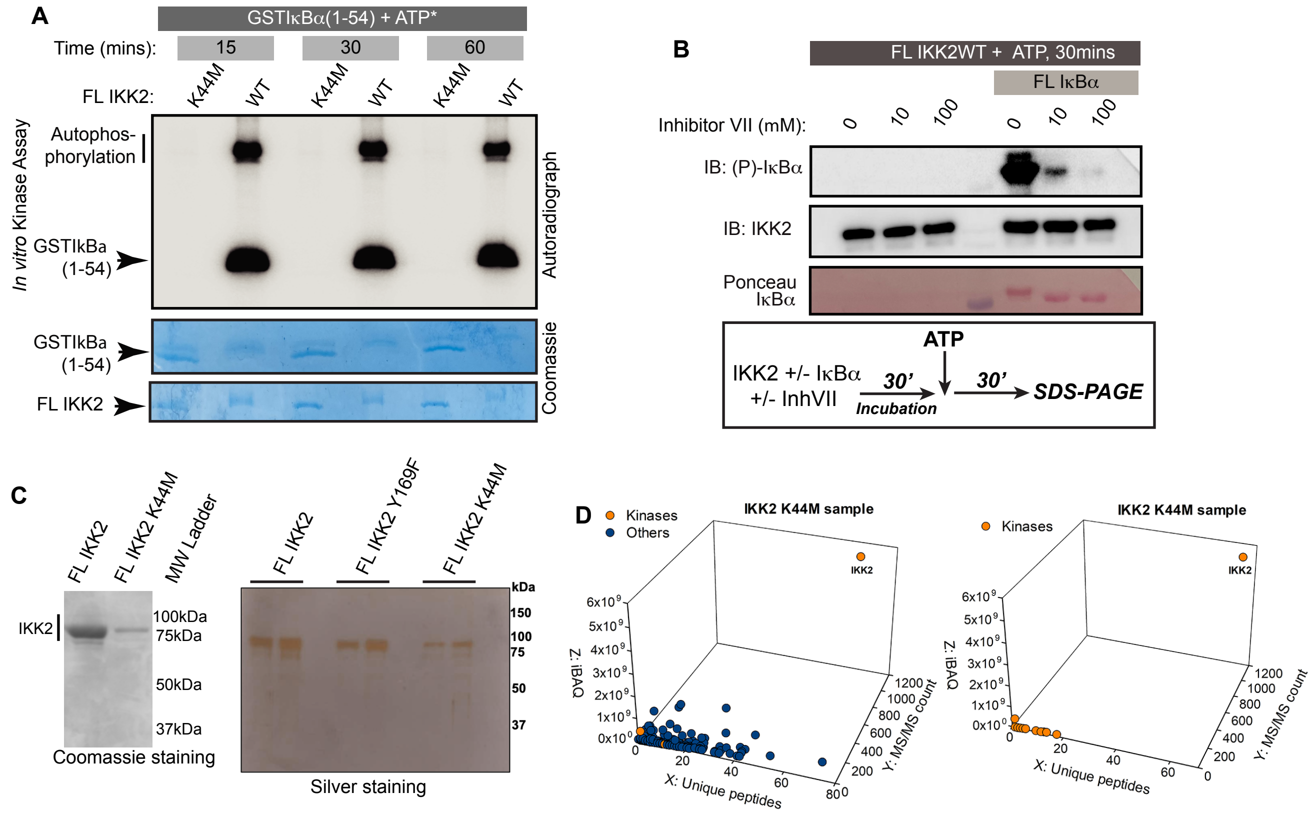

### Supplementary Figure 2

#### Figure 2 - Figure Supplement 1

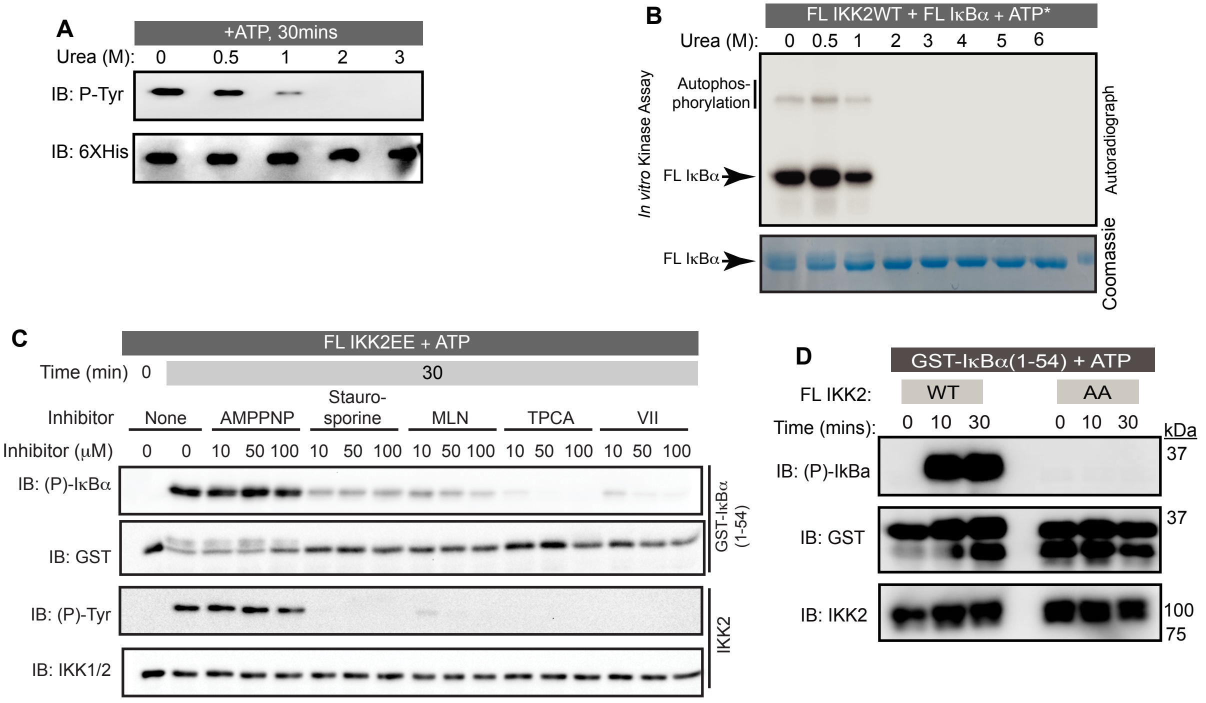

### Supplementary Figure 3

#### Figure 3 - Figure Supplement 1

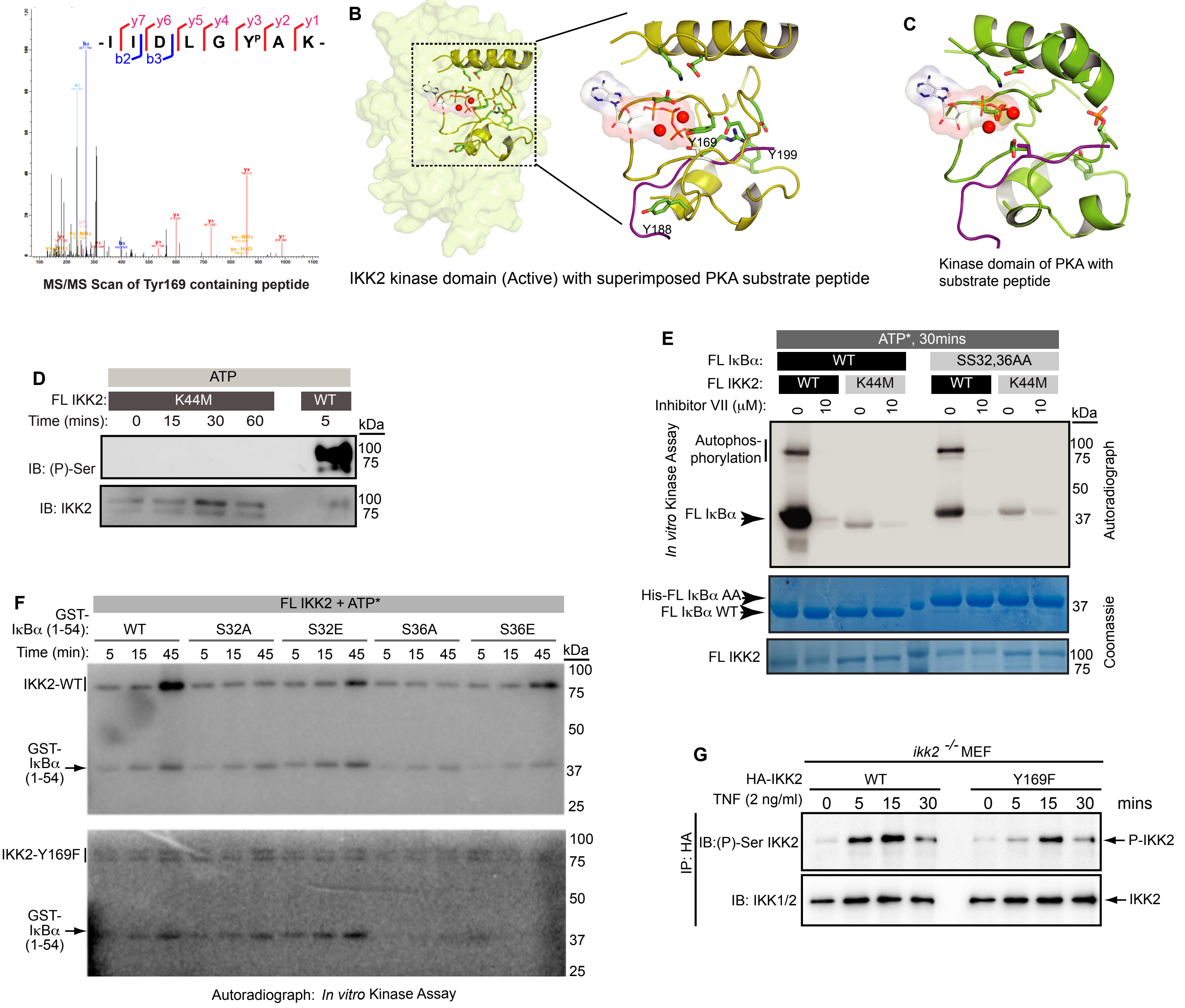

### Supplementary Figure 4

#### Figure 4 - Figure Supplement 1

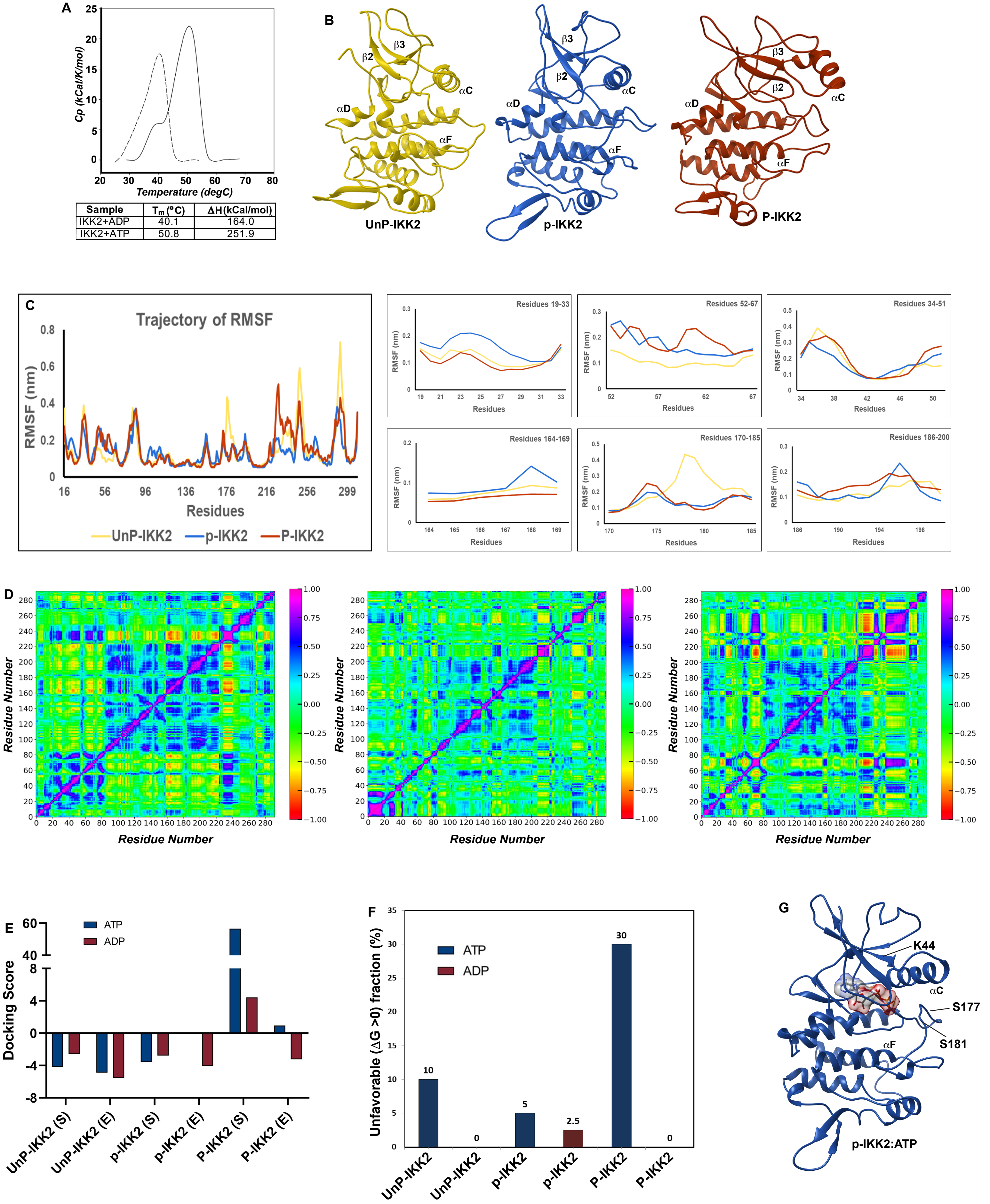


| Sample | $T_m$ (°C) | ΔH(kCal/mol) |
|---|---|---|
| IKK2+ADP | 40.1 | 164.0 |
| IKK2+ATP | 50.8 | 251.9 |

### Supplementary Figure 5

#### Figure 5 - Figure Supplement 1

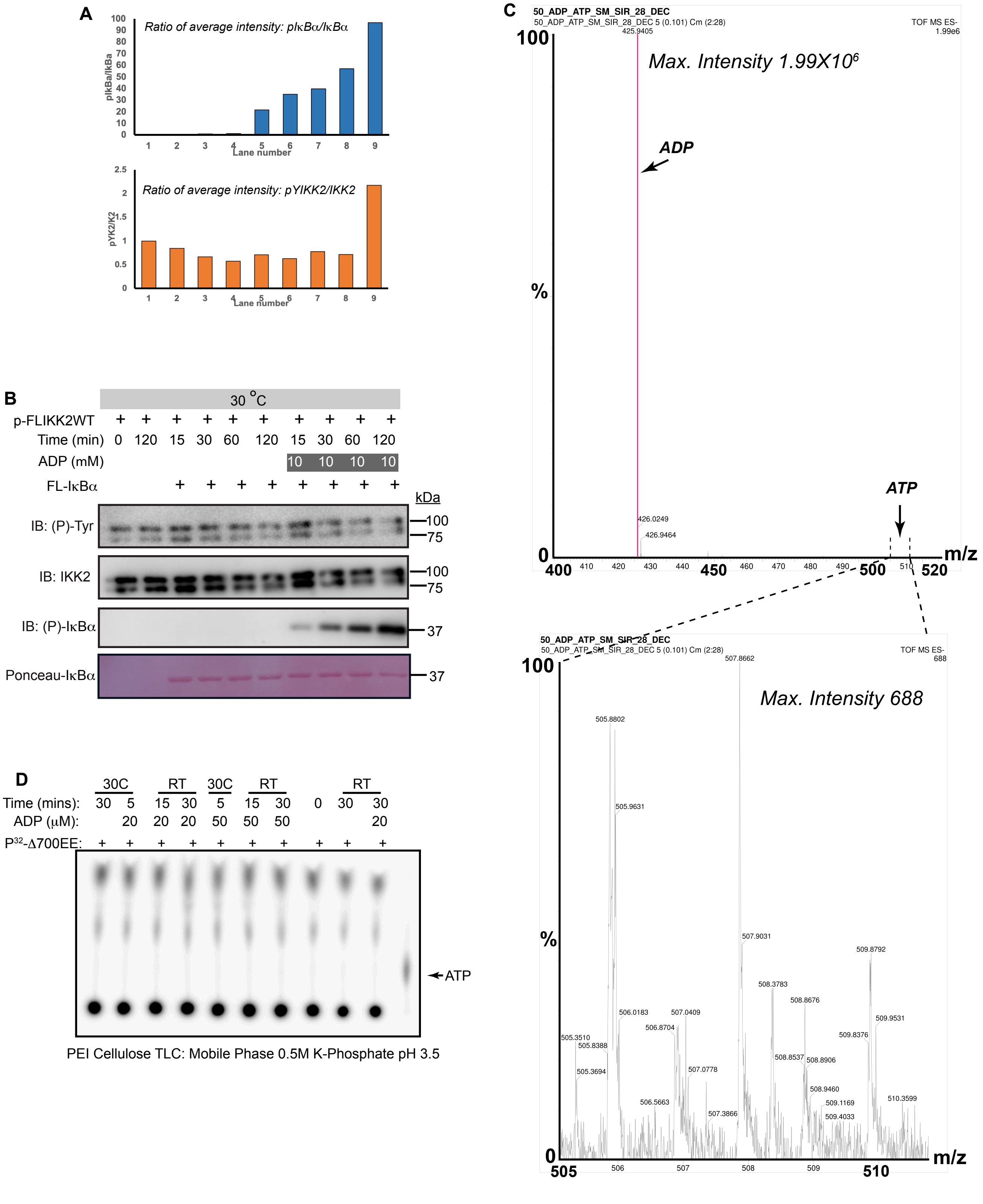

### Supplementary Figure 6

**Figure 6 - Figure Supplement 1**

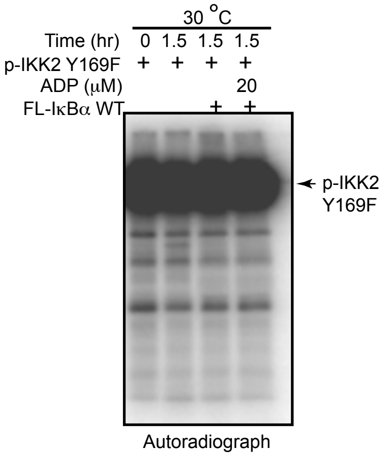
